# Different molecular signatures in lung cancer types from integrative bioinformatic analyses of RNASeq data

**DOI:** 10.1101/501569

**Authors:** Marta Lucchetta, Isabelle da Piedade, Mohamed Mounir, Marina Vabistsevits, Thilde Terkelsen, Elena Papaleo

## Abstract

**Background:** Genomic initiatives such as The Cancer Genome Atlas (TCGA) project contain data on profiling of thousands of tumors with different-omics approaches, providing a valuable source of information which may be used to decipher cancer signaling and related alterations. Managing and analyzing data from large-scale projects such as TCGA is a demanding task. Indeed, it is difficult to dissect the high complexity hidden in genomic data and to adequately account for tumor heterogeneity.

**Results:** In this study, we used a robust statistical framework along with the integration of diverse bioinformatic tools to analyze next-generation sequencing data from more than 1000 patient samples from two different lung cancer subtypes, i.e., the lung adenocarcinoma (LUAD) and the squamous cell carcinoma (LUSC). In particular, we used RNA-Seq gene expression data to identify both co-expression modules and differentially expressed genes to accurately discriminate between LUAD and LUSC. Moreover, we identified a group of genes which could act as specific oncogenes or tumor suppressor genes in one of the two lung cancer types, as well as two dual role genes. Our results have also been cross-validated against other transcriptomics data of lung cancer patients.

**Conclusions:** Our integrative approach allowed to identify two key features: a substantial up-regulation of genes involved in O-glycosylation of mucins in LUAD, and a compromised immune response in LUSC. The immune-profile associated with LUSC is linked to the activation of three specific oncogenic pathways which promote the evasion of antitumor immune response, providing new future directions for the design of target therapies.

## 1. Background

Lung cancer is one of the most aggressive cancers, with a five-year overall survival of 10– 15% [1]. Lung cancer can be classified into small cell lung cancer (SCLC) and non-SCLC (NSCLC), which account for 15% and 85% of all lung cancers, respectively. The main subtypes of NSCLC are divided mainly into adenocarcinoma (LUAD) and squamous cell carcinoma (LUSC). Lung cancer is a highly heterogeneous cancer type with multiple histologic subtypes and molecular phenotypes [2,3].

Since 2015, the classification of lung tumors has been defined by both their cytology and histology [1,4]. Despite the staining strategy to separate the lung tumors into different classes, cases that are immunohistochemically ambiguous are often reported and difficult to resolve. A proper differentiation between LUAD and LUSC also determines eligibility for certain types of therapeutic strategies [5]. For example, certain drugs are contraindicated for one of the two lung cancer types, such as Bevacizumab (Avastin) in LUSC [6]. It thus becomes crucial to discriminate among the two lung cancer types in a precise way. Microarray technologies have been used to identify differentially expressed genes (DEGs) in lung cancer samples to identify critical markers [7–10]. For example, Naval et al. identified a prognostic gene-expression signature of 11 genes which was subsequently validated in several independent NSCLC gene expression datasets[9]. This pioneering study established the prognostic impact of carcinoma-associated fibroblasts gene-expression changes in NSCLC patients. However, the markers identified in the study do not differentiate between LUAD and LUSC.

Gene or microRNA markers may be used, in principle, to distinguish between these two types of cancer [11–14]. Most of the studies carried out so far on LUAD and LUSC gene expression data have focused on the selection of a group of genes without considering, for example, the co-expression of the candidate genes. Single markers are very unlikely to be sufficiently robust to discriminate between cancer subtypes due to the intrinsic heterogeneity of tumors. In this context, new methods have been developed for robust analyses of co-expression signatures in gene expression data [15–17],

The Cancer Genome Atlas (TCGA) is a large genomic initiative in which more than 10 000 patients were profiled using six different platforms to identify cancer-related signatures [18–20]. TCGA provides a unique resource which can be re-analyzed for the discovery of cancer-related alterations or new biomarkers specific to a certain cancer (sub)type. Among the next-generation sequencing (NGS) platforms available, RNA-Seq is a recent and reliable approach for quantification of changes at the transcriptional level [21]. Lung cancer datasets for LUAD [22] and LUSC [23] are available in TCGA and account for more than 1000 samples overall. In parallel, the Recount2 initiative [24], which integrates GTEX [25] and TCGA, has recently allowed for an increase of healthy tissue samples for the comparison with tumor samples. Thus, the LUAD and LUSC TCGA datasets offer a suitable framework for the identification of gene expression signatures that could discriminate between the two lung cancer types regarding classification, diagnosis, and prognosis, as well as to shed light on the underlying molecular mechanisms. These two datasets have been used either to identify general cancer signatures [10,26–32] or to pinpoint signatures specific in only one of the lung cancer types [33–35].

Cline and colleagues [32] recently showed that there is a subset of 19 samples in the LUSC cohort that feature a LUAD-like gene expression profile. They labeled these samples ‘discordant LUSC’. Discordant LUSC samples are borderline for subtype classification, and the similarity with LUAD is also modest. These findings were also supported by the analyses from Piccolo’s group on an alternative pre-processing of the TCGA datasets [31]. As such, it is important to account for this information in the re-analysis of the TCGA lung cancer data to avoid misleading conclusions.

We aimed to closely compare LUAD and LUSC TCGA datasets using a robust statistical and bioinformatic framework. In particular, we have: i) identified a group of genes that are differentially expressed between LUAD and LUSC lung cancer types when compared to the normal samples, ii) assessed changes in the gene expression signature over cancer stages, iii) identified modules of differently co-expressed genes in the two lung cancer types and how the transcription regulation of the module genes was altered, iv) predicted potential oncogenes, tumor suppressors or dual role genes for each type and, iv) evaluated if there were potential prognostic markers among the group of LUAD- or LUSC-specific candidate genes. Overall, our study resulted in a subset of genes and pathways that have the potential to be used to discriminate among the two cancer types. Moreover, we identified candidate genes which deserve further functional/structural studies since they are poorly understood but potentially important as lung cancer markers or targets. Lastly, our data can also provide a useful guide to cellular studies using cancer cell lines which reflects the LUAD or LUSC types.

## 2. Results

### 2.1 Curation and description of the datasets used in the study

The datasets for our analyses were curated to remove the LUSC discordant samples and to remove samples with a low tumor purity (see Materials and Methods). The number of samples and genes retained for the analyses are reported in **Table SI.**

We performed differential expression analyses (DEAs) using generalized linear models to identify a subset of differentially expressed genes in the two TCGA lung cancer datasets LUAD and LUSC tumor primary (TP) with respect to the normal (NT) samples. A clear consensus on the best DEA approaches for RNA-Seq data does not exist yet, and different DEA methods could provide different results [36–40]. We thus employed three pipelines for DEA of LUAD and LUSC and generated a consensus list of DE genes (see Materials and Methods).

In addition, we curated three different datasets for each cancer type: i) two datasets containing all the samples (LUAD_all_, LUSC_all_) that account for partially paired tumors and normal samples; ii) two datasets containing only paired samples (LUAD_paired_, LUSC_paired_), i.e., normal and tumor samples from the same patient; iii) two datasets containing all samples without paired tumor samples (LUAD_unpaired_, LUSC_unpaired_). The pre-processed and processed (after normalization and filtering steps) data used in the study are deposited in our *Github* repository (https://github.com/ELELAB/LUADvsLUSC_tcga). A summary of the genes found to be up- or down-regulated by the different combination of curated datasets and DEA approaches is also reported in the *Github* repository. The *paired* datasets aimed to account for the proper removal of individual variability. We noticed that the usage of paired datasets from TCGA made a difference in the comparison of individual gene levels [41] and we thus aimed to evaluate its impact more broadly on the DEA analyses. The usage of the third dataset (*unpaired*) aimed to remove artifacts due to a partially paired dataset [42] with the purpose of identifying a gene expression profile that may be observed without correction for patient-specific effects. The choice to remove the paired tumor samples was dictated by the fact that the datasets contained several non-paired tumor samples, whereas almost all the normal samples were associated with a paired tumor sample and were thus retained. A comparison of the DE genes obtained from the analysis of each of the three datasets allowed us to evaluate the impact of different curations in terms either of either sample size or sample pairs.

### 2.2. The usage of paired or non/partially paired data mildly affects the results of DEA, whereas a too simplistic design in the DEA protocol has marked effects

At first, we assessed the influence of using different definitions of the dataset for DEA analyses. We compared the results of DEA carried out with a certain method, i.e., *limma* or *edgeR* or *edgeR-TCGAb*, on the three different datasets *{paired*, *all*, *unpaired)* for each of the two cancer types (LUAD and LUSC) (Figure 1). The differences in the total number of DE genes were minimal among the different datasets (approximately 1-8 %). We noticed that the usage of paired data provides a more stringent selection of up-regulated genes but is more permissive concerning the estimate of down-regulated genes. Moreover, the datasets *unpaired* and *all* feature a more substantial overlap with respect to the *paired* ones (Figure 1).

**Figure 1.**
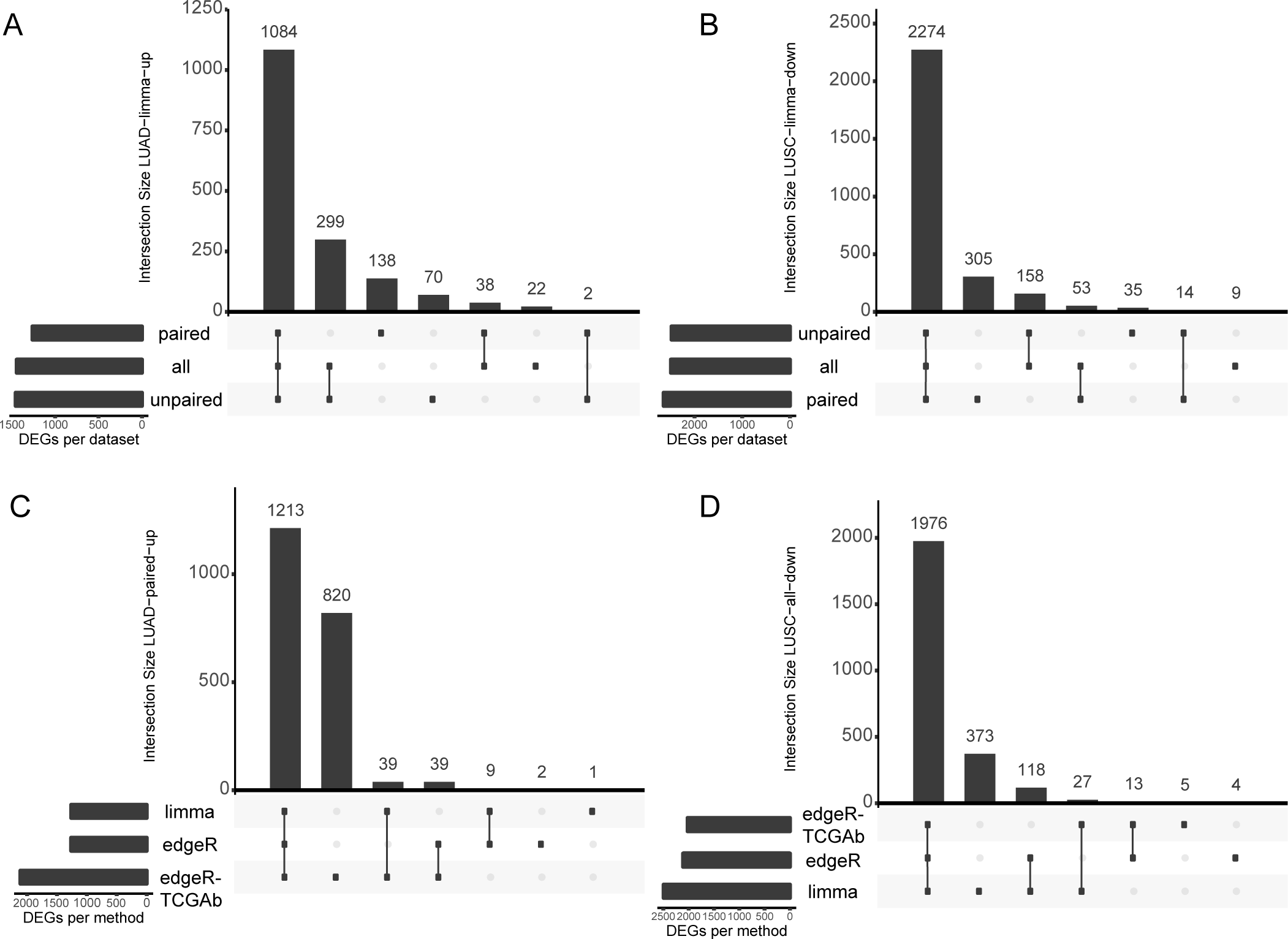
Comparison of DEA results with different curation of the datasets and different DEA protocols. For sake of clarity, we reported the example of up-regulated genes in LUAD (A,C) and down-regulated genes in LUSC (B,D) with the *limma* approach. The other approaches gave similar results and can be reproduced through the scripts in our Github repository (https://github.com/ELELAB/LUADvsLUSC_tcga).

Next, we analyzed in more detail, the impact of the different DEA approaches on the definition of DE genes. *Limma* results in the most stringent approach in the case of up-regulated genes **(Table S2).** Inversely, *limma* provides a large number of down-regulated genes (more than 300) that are not identified by the two *edgeR* pipelines (Figure 1). At the gene identity level, we also observed a similar pattern to the one above, i.e. that partially paired *(all)* and *unpaired* datasets have similar trends regarding the influence of the DEA approach.

We obtained the most pronounced differences in the case of paired data. In this case, *edgeR-TCGAb* featured a subset of up-regulated genes which are not identified by the other two methods (Figure 1). We noticed a similar effect, even if less pronounced, when unpaired or all samples were used (see data in our *Github* repository). In the case of paired samples, this behavior can be explained by the fact that the *edgeR-TCGAb* pipeline does not correct for patient-specific effects which are likely important when the normal and tumor samples are matched. In the case of partially paired or unpaired samples, the effect is less pronounced, and it is likely to be related to the fact that *edgeR-TCGAb* DEA pipeline does not include batch corrections which we included directly within the design matrix in the other two DEA pipelines. We therefore decided to have a closer look at the 820 and 619 up-regulated genes identified only by the *edgeR-TCGAb* pipeline in LUAD and LUSC, respectively. We extracted these genes and compared their logFC and FDR values, as calculated by the three approaches (Figure 2). Most of the discordant genes have either logFCs close to or below to 1 or FDR values close to 0.01 (50-60% of the genes), i.e., they are borderline significant according to the DEA criteria set for analysis with *edgeR* or *limma*. Moreover, *edgeR-TCGAb* tends to overestimate the logFC values. We also noticed that there is a small number of cases in which *edgeR-TCGAb* also assign an opposite directionality, i.e., the genes are down-regulated according to the other two methods. Specifically, this set of genes included: DUOXA2, IGFALS and, KLK14 in LUAD, as well as EDN3, GFI1B, MYH15, PEG3 and PENK in LUSC. We searched for each of these non-congruent genes in the *IGDB*.*NSCLC* database [43], which is a collection of genes that are known to be altered in NSCLC. We did not consider those hits with significant p-values but for which either the probe sets are reported with mapping problems or the fold change is lower than 2. We observed that only MYH15 is found to be up-regulated in one of the LUSC studies at *IGDB*.*NSCLC*, while the other genes listed above are significantly down-regulated, supporting the *edgeR* and *limma* results from our study.

**Figure 2.**
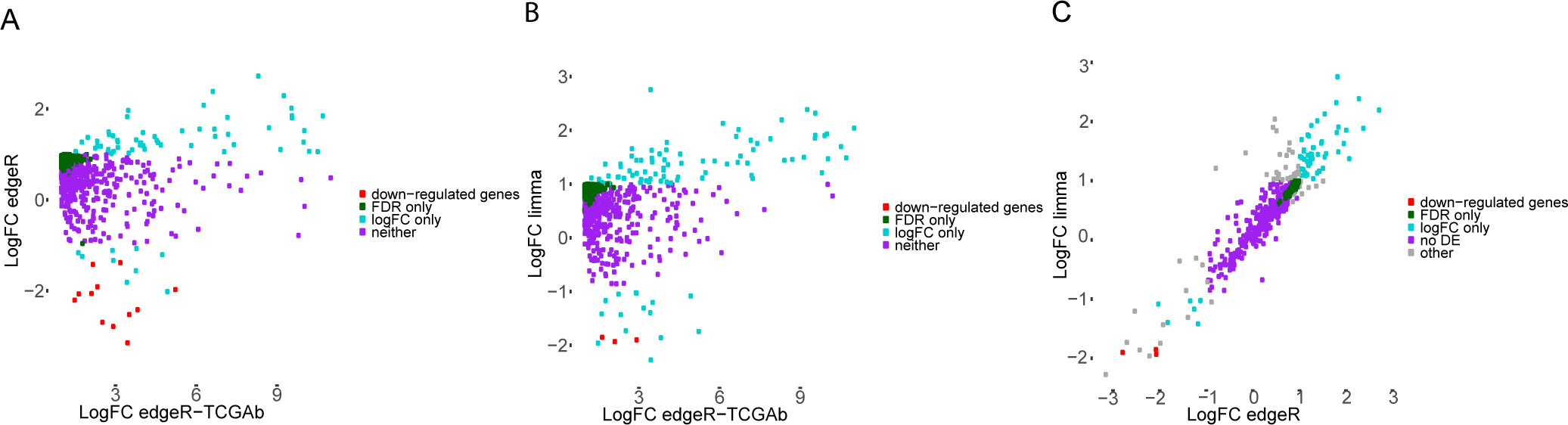
Analysis of the 820 up-regulated genes identified only *by edgeR-TCGAb* and gene enrichment analyses. The comparison between *edgeR-TCGAb* with *limma* and *edgeR*, respectively are shown in the upper panels, whereas the scattered plot comparing *limma* and *edgeR* are shown in bottom left panel. In the upper panels, we reported the genes that: i) are identified as down-regulated (in red), ii) have a significant FDR but not logFC (in green), iii) have a significant logFC but not FDR (in cyan), iv) are not significant according to both FDR and logFC (purple), according to *edgeR* (A) or *limma* (B) approaches. In the panel C, we used a similar color code, the only difference was that the condition is satisfied by both the *edgeR* and *limma* approaches (i.e. in red down-regulated genes for both the methods). Those genes that are in disagreement between *limma* and *edgeR* are shown in grey. We obtained the same results for LUAD and LUSC and we reported the LUAD case as an example.

The results of our analyses, thus, raise concerns about the accuracy of the original *TCGAbiolinks DEA* pipeline with *edgeR (edgeR-TCGAb)* especially when paired samples are analyzed, highlighting a need for a different DEA design within the R/Bioconductor package. This design should include functions for proper batch corrections and corrections for patient-specific effects, which we recently implemented in the current version of *TCGAbiolinks* (v 2.8).

We observed the main differences in DEA genes when the *paired* dataset was compared to the other two datasets, as expected, considering that we explicitly corrected for the patient-specific effects in the *paired* scenario. We also concluded that there is no significant advantage in the usage of unpaired samples in the DEA of TCGA datasets, at least when only a small fraction of paired tumor samples is removed (i.e., less than 10% of the whole tumor dataset). Overall, 60-80% of the DE genes are in common among the three methods, suggesting that their integration may allow for removal of genes with borderline significance with the purpose of defining a robust signature of LUAD- and LUSC-specific genes.

### 2.3 Identification of LUAD- and LUSC-specific differentially expressed genes

In the following steps, we employed the LUAD_all_ and LUSC_all_ datasets since they are the ones that allow us to maximize the sample size. The results of the other two dataset curations were mainly used as a further control in each of the analyses. Moreover, as an additional control of our analyses, we carried out DEA on the LUAD and LUSC re-processed *unified* datasets from the recent *Recount2* initiative [24]. In the *Recount2* platform, TCGA data were integrated with the normal GTEX (Genotype Tissue Expression Project)[25] samples (see Materials and Methods for details). This integration increases the pool of available normal samples for the comparison for a total of 374 healthy samples. Moreover, *Recount2* also provides a genuine source of healthy tissue samples to compare with the lung tumors, e.g., not only to the normal adjacent tissues that are available in the TCGA. The list of DE genes for LUAD and LUSC for the *unified* datasets are reported in our *GitHub* repository.

We employed a consensus approach, in which we defined as DE genes in LUAD and LUSC only those found by all the three DEA approaches (i.e., the intersects in each of the overlap diagrams similar to the ones reported in Figure 1, bottom panels) in the datasets *all*. We then compared the up- and down-regulated genes in LUAD_aii_ with the ones of LUSCaii. Indeed, to identify gene signatures that can differentiate between the two lung cancer types, it is not sufficient that the genes are differentially expressed with respect to the normal samples. We also need to verify that they are not up (or down) regulated in both the cancer types. Moreover, we further pruned the list of specifically up- (or down-) regulated genes for LUAD or LUSC by those genes that are identified as specifically up- (or down-) regulated in the opposite cancer type with the DEA consensus approach applied to the datasets *paired* to filter out false positives.

Overall, we retained 337 and 1451 genes specifically up-regulated, as well as 165 and 956 down-regulated genes in LUAD and LUSC, respectively (reported in our Github repository). Interestingly, we identified a small subset of genes that were up-regulated in LUAD but down-regulated in LUSC (MUC5B, HABP2, MUC21, and KCNK5) or vice-versa (CSTA, P2RY1, ANXA8, NELL2, and NTRK2).

We carried out pathway enrichment analysis using *ReactomePA* [44] for the up- and down-regulated unique genes of LUAD and LUSC. The analyses revealed that pathways related to O-linked glycosylation of mucins is enriched for the up-regulated genes of LUAD and down-regulated genes of LUSC, respectively (Table 1). This suggests that the proteins involved in this pathway could play an important role in discriminating between the two lung cancer types. Of particular interest is a group of mucins (MUC1, MUC4, MUC5B, MUC13, MUC15, MUC16, and MUC21), as well as enzymes involved in their modifications. Additionally, we noticed that genes involved in the complement system (C2, C3, C4BPA, C5, CFH, and CFI) and several genes related to the pathways of the innate immune response are significantly downregulated in LUSC. We obtained similar results using GO-enrichment analysis on GO biological processes on LUAD samples with regards to O-linked glycosylation **(Figure IS).**

**Table 1.** Pathway enrichment analysis with *ReactomePA*. Only the results relevant to the comparison of O-glycosylation, immune response and complement pathways are reported. For the full list of results, one could refer to our Github repository for the project.

| Pathway ID | Description | FDR | Gene IDs |
| --- | --- | --- | --- |
| 913709 | O-linked glycosylation of mucins | 0.01 | LUAD(up):<br>B3GNT6/GALNT6/MUC13/MUC16/MUC21/MUC4/MUC5B |
| 913709 | O-linked glycosylation of mucins | 0.04 | LUSC(down):<br>B3GNT7/B3GNT8/GALNT10/GALNT5/MUC1/MUC15/MUC21/MUC5B/<br>ST3GAL2/ST6GALNAC4 |
| 977068 | Termination of O-glycan biosynthesis | 0.01 | LUAD(up):MUC13/MUC16/MUC21/MUC4/MUC5B |
| 5173105 | O-linked glycosylation | 0.02 | LUAD(up):<br>B3GNT6/GALNT6/MUC13/MUC16/MUC21/MUC4/MUC5B |
| 5173105 | O-linked glycosylation | 0.02 | LUSC(down):<br>B3GNT7/B3GNT8/GALNT10/GALNT5/MUC1/MUC15/MUC21/MUC5B/SPON1/ST3GAL2/ST6GALNAC4/THBS1 |
| 977606 | Regulation of Complement cascade | 0.001 | LUSC(down):C2/C3/C4BPA/C5/CD55/CFH/CFI/CR1 |
| 168249 | Innate Immune System | 0.01 | LUSC(down):<br>ADCY6/ADCY7/BIRC3/C2/C3/C4BPA/C5/CASP10/CCR2/CCR6/CD14/CD3G/CD55/CD80/CFH/CFI/CR1/CYFIP2/DTX4/DUSP6/ELMO1/FCER1A/FCGR2A/FGF7/FYN/GAB2/GRAP2/HLA-C/ICAM3/ITK/ITPR1/ITPR2/JUN/KLB/LGALS3/LRRFIP1/MEF2C/MYD88/MYO1C/NFATC2/NFKBIA/NOD1/PDGFRB/PLCG2/PRKCD/RIPK3/RNF125/RPS6KA1/RPS6KA2/TEC/TLR2/TLR3/TMEM173/TREM1/TXNIP/UNC93B1/VAV1/WIPF1 |
| 166658 | Complement cascade | 0.02 | LUSC(down): C2/C3/C4BPA/C5/CD55/CFH/CFI/CR1 |

### 2.4 Clustering genes in LUAD and LUSC across tumor stages

We aimed to identify a subset of specific and interrelated genes which, as an ensemble, could be more effective than single markers in discriminating between LUAD and LUSC. For this purpose, DEA alone is not sufficient. We therefore analyzed the molecular signatures both using soft-clustering approaches over the cancer clinical stages and implementing weighted co-expression analyses (see 2.5).

We applied a soft-clustering approach [45,46] to separate LUAD and LUSC genes into clusters based on their changes in gene expression during different cancer clinical stages [47], allowing us to identify six main clusters with different signatures over stages (Figure 3 and Figure S2). We observed that LUSC genes selected for each cluster have a higher membership value (Figure 3), i.e., more solid classification within the cluster, whereas the LUAD gene had a ‘soft’ membership **(Figure S2).**

**Figure 3.**
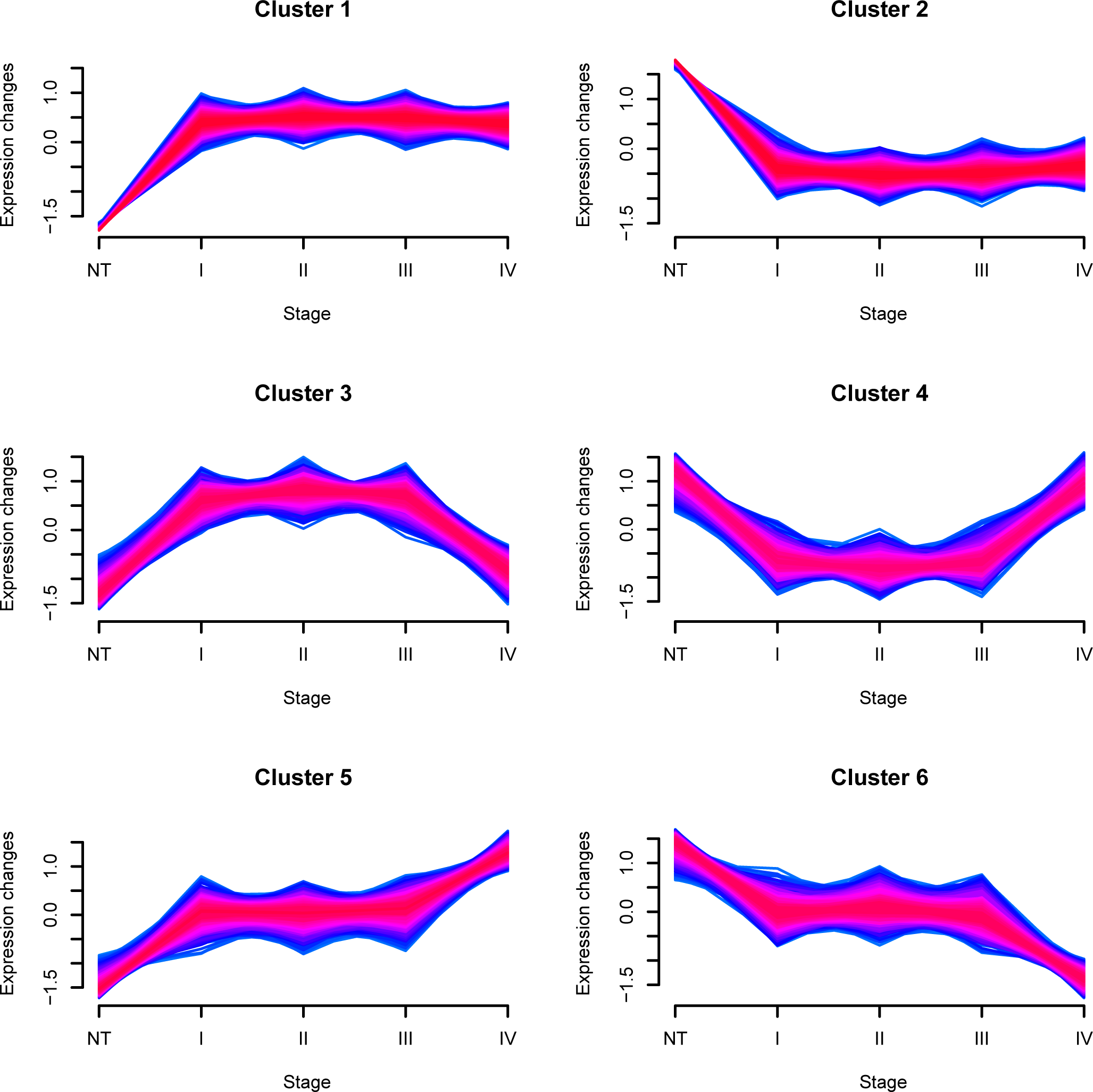
Soft-clustering across lung cancer clinical stages. Each cluster describes an expression pattern in the dataset through the four stages of cancer i.e stages I, II, III and IV. Blue and purple lines correspond to genes with high cluster membership value, *i*.*e*. m> 0.56. A Table with the genes belonging to each cluster and their m value is reported in the Github repository. The example of LUSCaii is showed for sake of clarity and the results for LUAD_all_ are reported in Figure S2.

Clusters 1 and 5 of LUSC (Figure 3), as well as clusters 1 and **6** of LUAD show a general up-regulation of genes along all stages **(Figure S2).** In contrast, clusters 2 and **6** of LUSC (Figure 3), and clusters 2 and **4** of LUAD **(Figure S2)** feature a general down-regulation when the four clinical stages are compared to the normal samples. We extracted the genes that are up-regulated in a certain cancer type and down-regulated in the other or the opposite, similarly to what we previously did in the context of the DEA results. We identify a group of **46** genes which are up-regulated in LUAD but down-regulated in LUSC. 72 genes are identified as down-regulated in LUAD and up-regulated in LUSC. The soft-clustering comparison thus provided an additional list of interesting gene candidates of which MUC5B, CSTA, P2RY1 and NTRK2 were shared between the soft-clustering and the DEA.

Clusters 3 of both LUAD and LUSC (Figure 3 and Figure S2) feature a signature in which the genes are up-regulated at the early clinical stages, but they decrease again at late stages (i.e. stage IV), whereas cluster 4 of LUSC and 5 of LUAD show the opposite trend, i.e. a down-regulation at early stages but increase at late stages (Figure 3 and Figure S2). These patterns may be indicative of dual-role/moonlight genes.

Expression of dual-role genes may, for example, be unwanted by cancer cells in early tumor stages, whereas they become essential later on in tumorigenesis, providing the cancer cell with a functional advantage or resistance to chemotherapy [30].

### 2.5 Prediction of oncogene and tumor suppressor genes in LUAD and LUSC

Genes that are up- or down-regulated and are also known to be oncogenes or tumor suppressors, respectively, are of great interest in cancer.

We therefore carried out a prediction of potential tumor suppressor genes (TSGs) and oncogenes (OGs) using the *Moonlight* workflow[30], which employs gene expression signatures and biological pathways to identify potential TSGs and OGs. This analysis is useful to integrate and expand the information available on TSGs and OGs through the curated data from *TSGene (TSGDB)[*48], *ONGene* [49] and *COSMIC* [50] (see Materials and Methods for details).

At first, we were interested in evaluating which of the up- and down-regulated genes that discriminate between LUAD and LUSC are known or predicted to be OGs (up-regulated genes) or TSGs (down-regulated genes).

We thus identified 24 potential TSGs and 146 OCGs for LUAD, while we obtained 22 TSGs and 456 OCGs for LUSC with the *Moonlight* approach (the details and full list of genes are reported in our *GitHub* repository). Only 31 predicted OCGs are common between LUAD and LUSC, whereas no TSGs are common among the two lung cancer subtypes. Intriguingly, IL6 and KRT23 are predicted as OCGs in LUAD but TSGs in LUSC, highlighting that these genes are deserving of attention in future studies. IL6 is of interest thanks to its role in the immune response and the complement system [51], as well as considering that it is down-regulated in LUSC. We also recently identified IL6 as the only down-regulated cytokine in breast cancer samples using cytokine assays [52]. However, IL6 downregulation could be also associated with the chemotherapeutic treatment as previously reported [53]. Future studies on naive tumor samples, as well as LUSC and LUAD cellular models where the IL6 gene could be overexpressed or silenced could shed light on its involvement in the two lung cancer types.

### 2.6 Coexpression signatures in LUAD and LUSC

As stated above, we seek gene expression signatures which may be useful in discriminating between LUAD and LUSC types, as well as interesting targets for each of the cancer types. For this reason, we carried out a gene co-expression analysis to identify different modular gene co-expression networks in LUAD and LUSC.

In LUSC, we identified six modules (Figure 4). Ml is enriched in proteins for organization and assembly of the cell and gap junctions, including gap junction proteins, like the up-regulated hub proteins GJB5, keratin type II proteins and protein channels activated by chloride. M2 is enriched in proteins for glutathione conjugation and response to redox stress, such as the up-regulated hub proteins sulfiredoxin-1 protein and the oxidative stress-induced growth inhibitor OSGIN1. M3 includes extracellular matrix organization and collage-related proteins. M4 is enriched in interferon signaling, cytokine signaling in immune response, with a down-regulation of HLA genes. M5 has no significant association to any annotated cellular pathway, whereas M6 is enriched in proteins that regulate the complement cascade.

**Figure 4.**
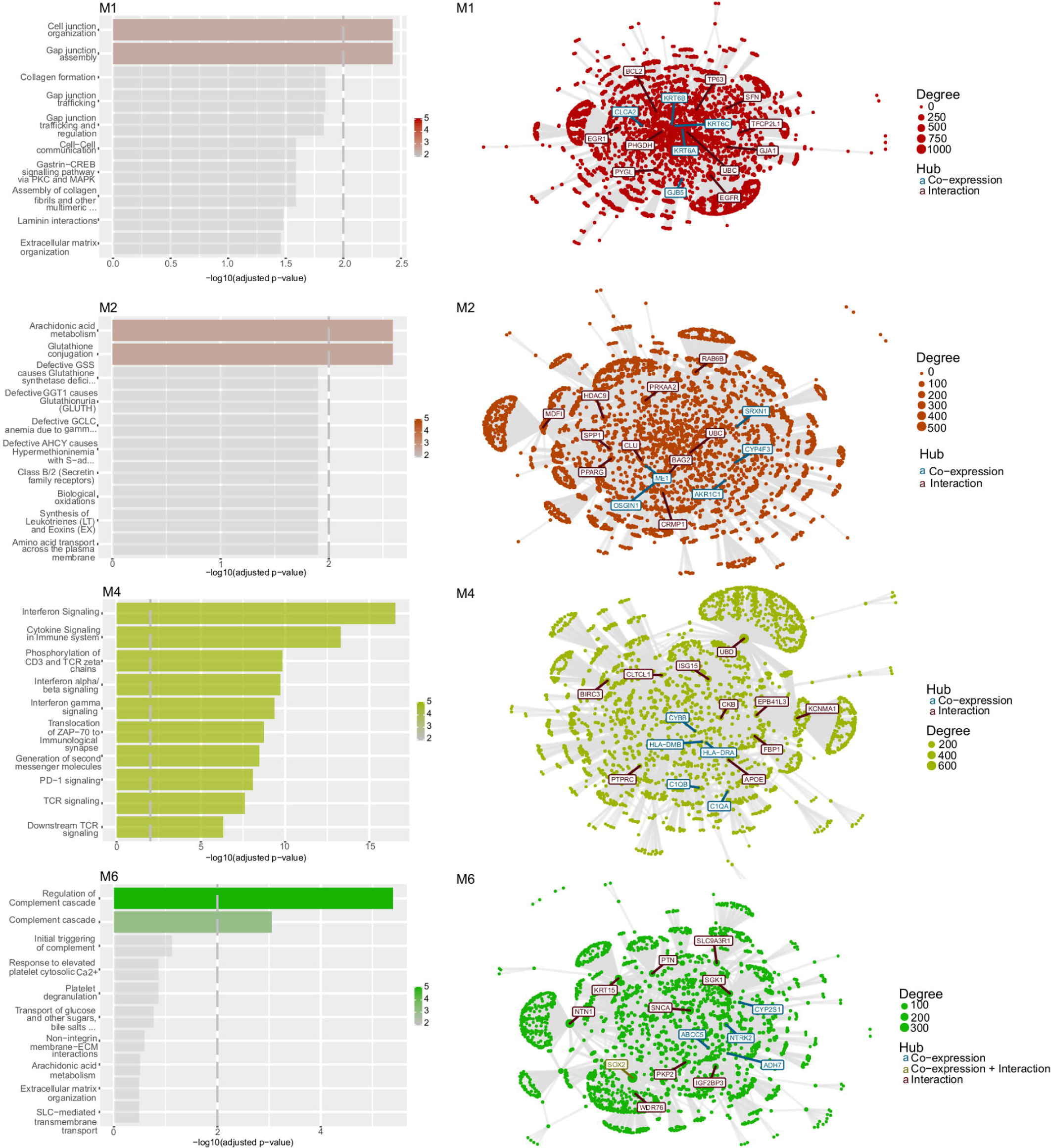
Coexpression modules and their network in LUSC. The modules which collect genes and pathways that differentiate LUSC from LUAD are shown in the figure, along with their networks built integrating the coexpression data with annotated protein-protein interactions.

We then identified four modules in LUAD (Figure 5): Ml which is enriched in extracellular matrix organization proteins and regulation of complement cascade; M2, which is enriched in interferon and cytokine signaling and, M3 which includes collagen-related genes and proteins for extracellular matrix organization. Notably, M3 is the only module that includes hubs which are conserved among LUAD and LUSC and thus not relevant to our study. M4 of LUAD has no significant associations with any known pathway.

**Figure 5.**
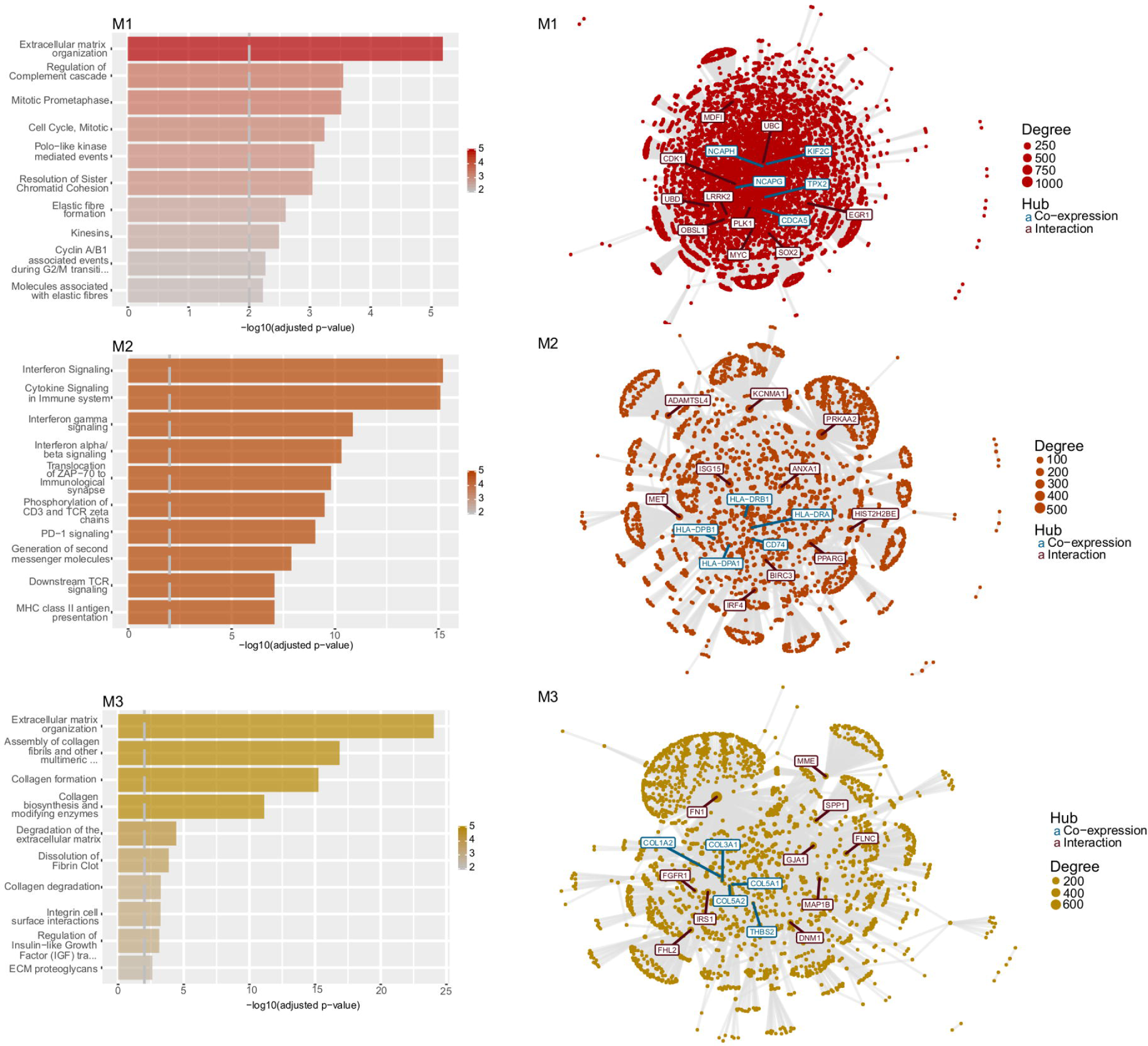
Coexpression modules and their network in LUAD. The modules which collect genes and pathways that differentiate LUAD from LUSC are shown in the figure, along with their networks built integrating the coexpression data with annotated protein-protein interactions.

Even if some of the modules of LUAD and LUSC are enriched for the same processes, a pairwise comparison of each of them suggested that, in most of the cases, the number of overlapping genes in the LUAD and LUSC modules is only a minor component. This could suggest that the genes triggering different pathways have different coexpression signatures in the two cancer types. Pathway-enrichment analyses on the DE and soft-clustering genes also pointed out a down-regulation of proteins involved in the complement cascade and genes related to the immune response in the LUSC samples (Table 1), enforcing the notion of a compromised immune response in LUSC.

Moreover, the Ml and M2 of LUSC are enriched in pathways that have not been found for the LUAD coexpression modules, i.e., pathways related to cellular junctions (Ml) and glutathione (M2).

For further analyses, we retained only those genes within each module that are truly unique for LUAD or LUSC, comparing each module of a cancer type to all the modules of the other cancer type. For each module, we extracted the known transcription factors and their targets using the *TRRUST* database as a source of information[54]. We identified a network of transcription factors and their targets for modules 1 and 2 of LUAD, as well as 1, 2,3 and 4 or LUSC (Figure 6). Out of these, LEF1 is of interest since it activates NRCAM in module 2 of LUSC - two genes that are also up-regulated in LUSC only. We also noticed the presence of the up-regulated CSTA in module 1 of LUSC which is transcriptionally regulated by FOS along with TP63 and its target gene ZNF750. In module 1 of LUAD, we identified an interesting network between the up-regulated gene AGR2 and its transcription factor FOXA1, along with the activator SPDEF.

**Figure 6.**
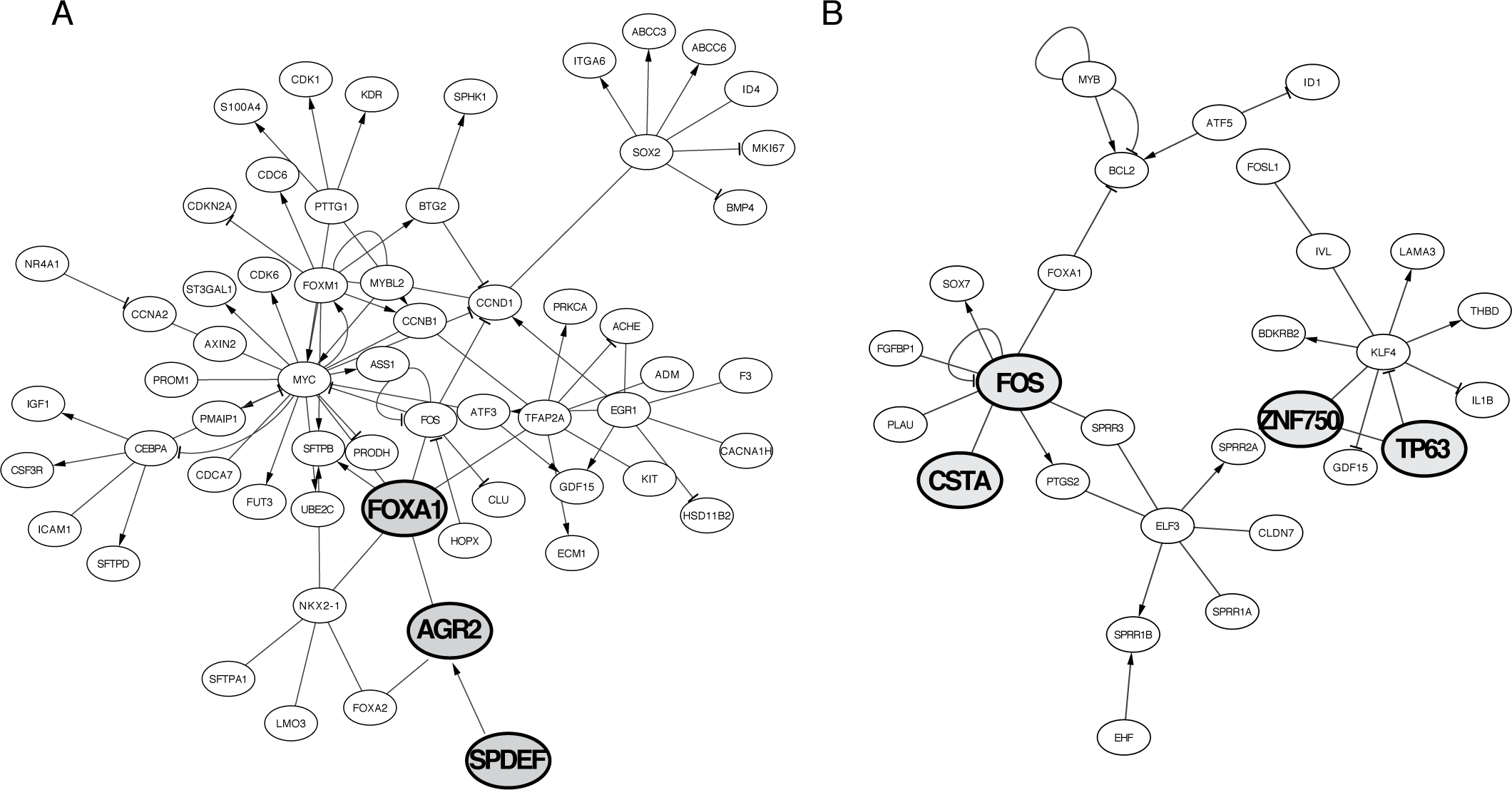
**Network of transcription factors and their target genes** in the coexpressed module 1 of LUAD (A) or LUSC (B).

### 2.7 Selection of candidate genes and their pathways

We collectively considered the results of the analyses described above with the final goal of proposing a subset of LUAD and LUSC-specific genes for further studies as markers or potential targets. In particular, we decided to retain only the genes that satisfy the following criteria: i) genes that are up- or down-regulated in a specific cancer type and the opposite in the other cancer type according to DEA and/or soft-clustering analyses, and ii) genes that belong to the coexpression modules with a functional signature and are truly unique to LUAD or LUSC. For each of these genes, we also annotated information on their potential as oncogenes, tumor suppressors or dual role genes; known associations with cancer according to the repository of disease-gene associations from text mining of the literature, *DISEASES* [55]. Specifically, we noticed if they matched with known oncogenes or tumor suppressors through analyses of *COSMIC* TGs and OCGs collection [50], *TSGDB* [48], *ONGene* [49] or prediction with the *MoonlightR* workflow [30] (see Section 2.5 and Materials and Methods). For dual role genes, we integrated as a reference for our study the curation from *TSGDB* [48], *COSMIC* [50] and the recently predicted ‘double-agent’ genes, namely Proto-Oncogenes with Tumor-Suppressor Functions (*POTSF*) [56]. This integrative reference annotation for dual role genes is reported in **Table S3** and in the *Github* repository for a total of 152 genes of which only 14 were all reported in all the three studies. The summary of each of the candidate genes along with their annotations is reported in Table 2.

**Table 2.** Candidate genes to discriminate between LUAD and LUSC in terms of gene expression levels, functions or prognosis. ‘N.S.’ indicates not significant results. The DisGeNET score and the DISEASES Z-score are provided in the table. *, **, and *** indicate a significant DE when the unified recount, the TCGA pair dataset or both are used.

| GENE | UNIPROT ID | DEA | Mfuzz cluster | CEMiTool Module | DISEASES | OG | TSG | DUAL ROLE |
| --- | --- | --- | --- | --- | --- | --- | --- | --- |
| MUC5B | Q9HC84 | Up(LUAD)<br>Down(LUSC) | Cl 6(LUAD)<br>Cl 2(LUSC) | M1(LUAD) | None | None | None | None |
| HABP2 | Q14520 | Up(LUAD)<br>Down(LUSC) | Cl 3(LUAD)<br>Cl 4(LUSC) | M1(LUAD) | 3.8 | None | None | None |
| MUC21 | Q5SSG8 | Up(LUAD)<br>Down(LUSC) | Cl 2(LUSC) | M1(LUAD) | 4.0 | None | None | None |
| KCNK5 | O95279 | Up(LUAD)<br>Down(LUSC) | Cl 3(LUAD)<br>Cl 2(LUSC) | M1(LUAD) | None | None | None | None |
| ICA1 | Q05084 | Up(LUAD)<br>Down(LUSC)* | Cl 6(LUAD)<br>Cl 2(LUSC) | M1(LUSC) | 1.9 | None | None | None |
| CSTA | P01040 | Down(LUAD)<br>Up(LUSC) | Cl 4(LUAD)<br>Cl 1(LUSC) | M1(LUSC) | 3.5 | None | TSGDB(CST5,<br>CST6) | None |
| P2RY1 | P47900 | Down(LUAD)<br>Up(LUSC) | Cl 4(LUAD)<br>Cl 1(LUSC) | M1(LUSC) | 2.2 | Moonlight(<br>LUSC,<br>P2RY14)<br>COSMIC(P<br>2RY8) | None | None |
| ANXA8 | P13928 | Down(LUAD)<br>Up(LUSC) | Cl 5(LUSC) | M1(LUSC) | 2.4 | None | TSGDB(ANXA1,<br>ANXA7) | None |
| FZD7 | O75084 | Up(LUSC) | Cl 4(LUAD)<br>Cl 1(LUSC) | M2(LUSC) | 3.7 | ONGENE(F<br>ZD2)<br>Moonlight(<br>LUSC,<br>FZD4) | None | None |
| ITGA6 | P23229 | Up(LUSC)*** | Cl 4(LUAD)<br>Cl 1(LUSC) | M1(LUAD) | 4.2 | ONGENE(I<br>TGA3)<br>Moonlight(<br>LUSC,<br>ITGA8) | TSGDB(ITGA5,IT<br>GA7, ITGAV) | None |
| CHST7 | Q9NS84 | Up(LUSC)** | Cl 4(LUAD)<br>Cl 1(LUSC) | M2(LUSC) | 1.6 | None | TSGDB(CHST10) | None |
| RND3 |  | Up(LUSC)* | Cl 4(LUAD)<br>Cl 1(LUSC) | M1(LUAD) | 3.3 | Moonlight(<br>LUSC,<br>RDN1) | TSGDB | None |
| ACOX2 |  | Down(LUSC) | Cl 2(LUSC) | M1(LUAD) | 2.2 | None | None | None |
| ALDOC |  | Up(LUSC) | Cl 1(LUSC) | M1(LUAD) | 3.9 | None | None | None |
| AQP5 |  | Down(LUSC) | Cl 4(LUSC) | M1(LUAD) | None | None | None | None |
| ARSE |  | Up (LUAD)*<br>Down(LUSC) | Cl 2(LUSC) | M1(LUAD) | None | None | None | None |
| FABP5 |  | Down(LUAD)<br>Up(LUSC) | Cl 4(LUAD)<br>Cl 5(LUSC) | M1(LUSC) | None | Moonlight<br>L(LUAD,FA<br>BP7) | TSGDB(FABP3) | None |
| SIPA1L2 |  | Down(LUAD)* | Cl 2(LUAD)<br>Cl 1(LUSC) | M3(LUAD) | None | None | None | None |
| SLC1A3 |  | Down(LUAD)* | Cl 2(LUAD) | M3(LUSC) | 3.5 | None | None | None |
|  |  |  | CI5(LUSC) |  |  |  |  |  |
| **SLC2A9** |  | Down(LUAD)* | CI4(DOWN<br>)<br>CI5(LUSC) | M1(LUSC) | None | None | None | None |
| **NRCAM** |  | Down(LUAD)*<br>Up(LUSC) | CI1(LUSC) | M2(LUSC) | 2.6 | Moonlight (LUSC, OCG) | TSGDB | None |
| **AGR2** |  | Up(LUAD)<br>Down(LUSC)** | CI3(LUAD) | M1(LUAD) | 4.1 | None | None | None |
| **SPDEF** |  | Up(LUAD)<br>Down(LUSC)** | CI1(LUAD)<br>CI2(LUSC) | M1(LUAD) | 3.6 | None | None | None |

### 2.8 Association of the gene signatures with patient survival

Next, we aimed to evaluate if any of the candidate genes had also a potential prognostic impact. We therefore carried out survival analyses using a Cox proportional hazard regression using all the candidate genes. We accounted for different explanatory variables, including the clinical stages, age and sex of the patients.

We retained only 14 and 16 genes out of the original list of LUAD and LUSC candidate genes, respectively since the others do not satisfy the proportional hazard assumption. Only four genes for LUAD (ITAG6, HABP2, FAB5P5 and RND3) are predictive for overall survival based on either FDRs and/or p-values **(Table S4)** regarding overall survival. In LUSC, we could not identify any significant prognostic marker for which the cox coefficient is in agreement with the direction of deregulation of the gene observed by DEA.

### 2.9 Cross-validation of the candidate genes

To further strengthen our results, we used two independent datasets (see Materials and Methods) where LUAD and LUSC samples were profiled by transcriptomics techniques with the same experimental setup as independent datasets for validation of the most interesting markers.

We retained only a subset of candidate genes reported in Table 2 for which LUAD and LUSC upper and lower quartiles are sufficiently separated when compared for the same gene so that they may suggest a potential value as a marker for classification of the two lung cancer types. We then extracted the ones for which gene quantification was available in the cross-validation datasets, and we used unsupervised clustering to verify if they can separate LUAD and LUSC. The results are reported in Figure 7. We could not validate the clustering potential of mucins since the data for these genes were not originally available as probes or after the probe set collapse operation. SPDEF, ICA1, FZD7, CHST7, SLC2A9, ACOX2, KCNK5, ARSE, P2RY1, RND3, CSTA, ALDOC, and ANXA8 from the coexpression modules 1 and 2 of LUAD or LUSC turned out to be promising for the separation of the two lung cancer types. KCNK5 was also proposed in a previous study [10]. We even identified a small core group of genes that still retain the capability of separating the two lung cancer subtypes, i.e., ALDOC, ARSE, ANXA8, and CSTA (Figure 7).

**Figure 7.**
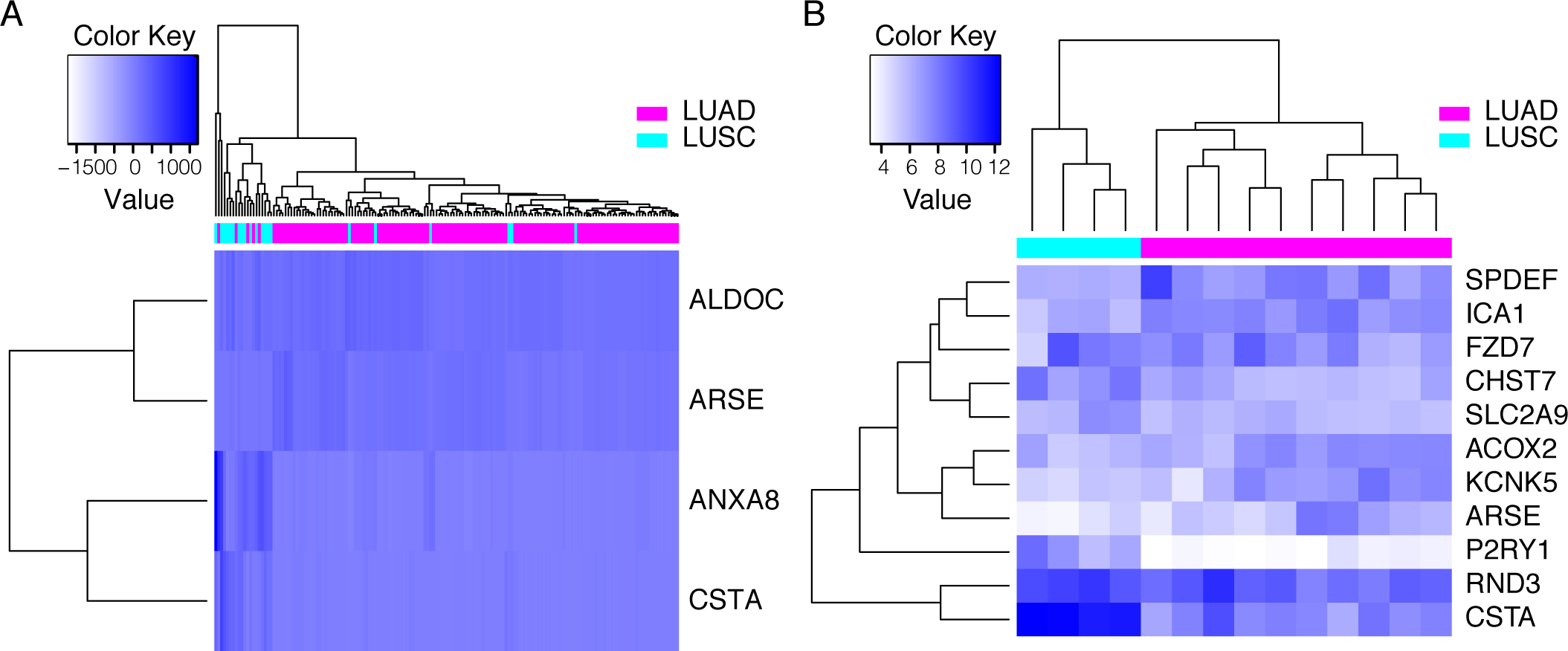
In silico validation of the candidate genes using other transcriptomics data for lung cancer. A minimal group of gene (core genes, panel A) or a more extended gene list (panel B) is showed that allow to classify the two cancer types.

## 3. Discussion

### Genes involved in O-glycosylation of mucins are differentially regulated in different lung cancer types

Our analyses pinpointed a differential regulation of different genes involved in the 0-glycosylation of mucins. These genes are up-regulated in LUAD and down-regulated in LUSC. Mucins are heavily glycosylated proteins where glycosylation is relevant to their function. Under normal conditions, mucins serve as a protective barrier for epithelial lung cells [57]. When dysregulated, these proteins promote cancer progression and metastasis [58]. During cancer progression, mucins can alone or in combination with different tyrosine kinase receptors mediate cell signals for growth and survival of cancer cells. Expression of certain mucins, such as MUC1 or MUC4 (identified also in our study), have been already associated to lung cancer in other studies, and in some cases even associated with poor prognosis for the patients [58]. Due to this key role in oncogenesis, mucins are emerging as attractive targets for novel therapeutic approaches to treat lung cancer and strategies have been already proposed [58].

Our results suggest that both membrane-bound (such as MUC21) and secreted mucins (such as MUC5B) contributes to the differences between LUAD and LUSC.

MUC5B overexpressing cancers more often show tendencies for relapse or metastasize postoperatively in comparison to non-expressing tumors [59], suggesting that LUAD patients could suffer from these events more often than the ones with a LUSC lung cancer type. Mucins are amenable drug targets, as attested by MUC1 which can be targeted by immunotherapy thanks to the availability of T-cell specific antigenic epitopes. Vaccines have been also proposed, as well as aptamer-based drugs (for a review [58]). Despite several studies on mucins in lung cancer, these have only scraped the surface of a complex and intricate interplay where also the interactions between the different mucins can add an extra level of undisclosed complexity. Our data suggest that more studies focusing specifically on MUC5B and MUC21 are needed, considering both the opposite behavior of these two proteins in LUAD and LUSC and overexpression in LUAD, which suggest the possibility of exploiting them (or the enzymes regulating mucin glycosylation) as drug targets for LUAD-specific therapy.

### LUSC and the activation of oncogenic pathways for evasion of antitumor activity

Generally, an enhanced immune response in cancer can be exploited for therapeutic purposes[60]. We here observed that LUSC seems to be immune-compromised with a signature of massive down-regulation of the complement cascade and other key genes for native immune response. Our results nicely fit within the recent scenario of overall difference in tumor immune landscape in LUAD and LUSC [61].

Recently, five main oncogenic pathways have been reported [62] that are associated with the evasion of antitumor immunotherapy. The activation of these pathways which relies, in turn, on the dysregulation or mutations of usual suspects in cancer such as p53, cMYC, and the P-catenin/WNT. These genes are the upstream regulators of these evasion pathways and act through a well-orchestrated cascade of other more specifically deregulated and diverse set of genes (Figure 8). The oncogenic pathways for evasion of immune response in tumor cells have the ultimate effect of impairing the induction or execution of a local antitumor immune response, which also explains the resistance to certain therapies. Using our integrative analyses on DE and coexpressed genes we were able to link the activation of three of these pathways to LUSC, providing new horizons for the design of new tailored therapies to this cancer type.

**Figure 8.**
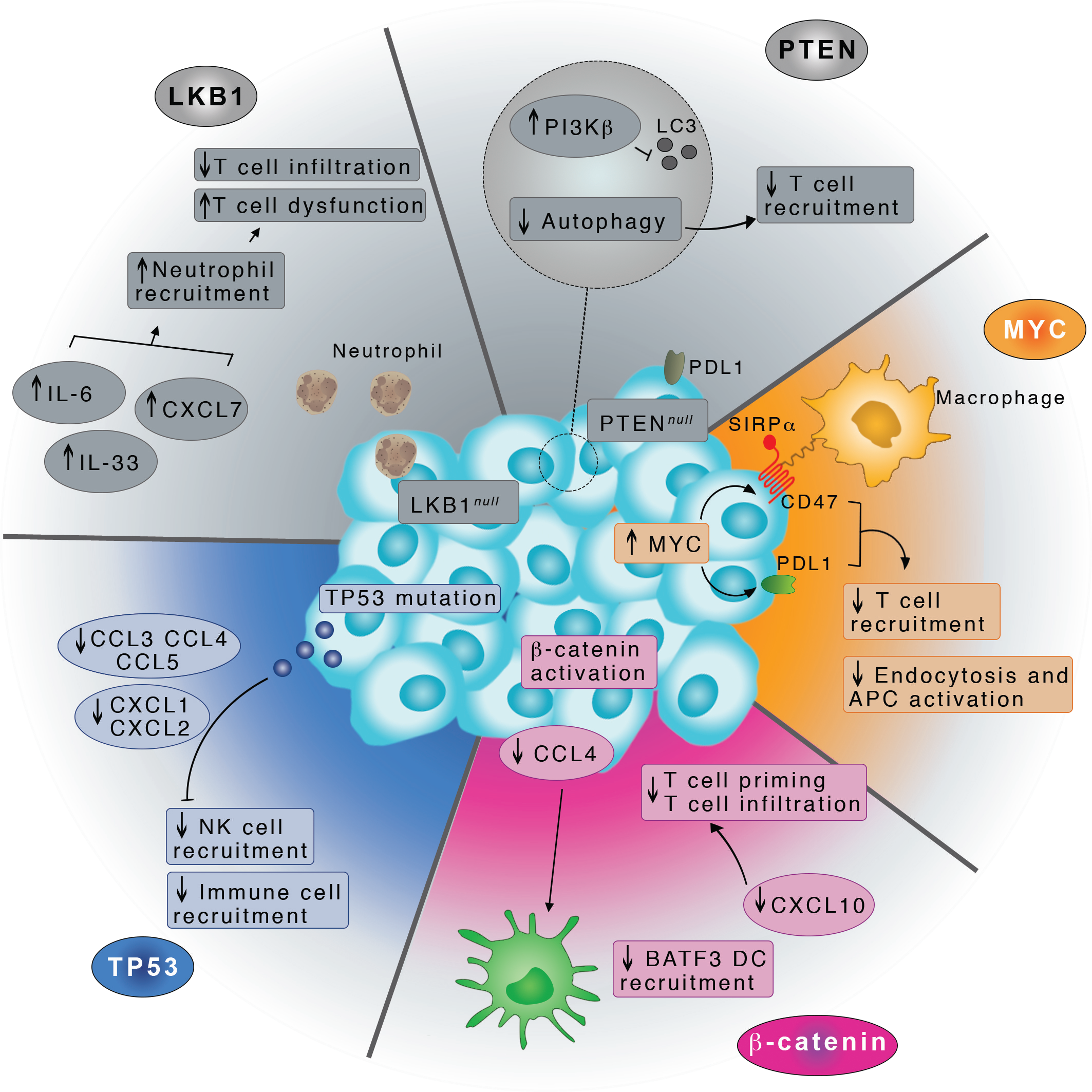
Illustration of the three oncogenic pathways to evade tumor immune response which we found activated in LUSC.

## 4. Conclusions

This study allowed to shed new light on the differences between two elusive lung cancer types, i.e., LUAD and LUSC. In addition, we here provide a useful integrative biostatistics and bioinformatics framework for the interpretation of gene expression data. Our results suggest that to use partially paired, paired or unpaired samples will not yield markedly different outcome from downstream analyses, e.g., differential expression or enrichment analyses. On the contrary, the protocol used for the DEA, especially in the context of the up-regulated genes and paired data, needs to be carefully assessed since too simplistic approaches without the proper information incorporated in the design matrix can results in discordant signatures.

We predicted two potential dual role genes (IL6 and KRT23) in LUAD and LUSC. Our analysis also showed that LUAD and LUSC differentiate for the biological processes that are altered. In particular, LUAD feature an up-regulation of genes involved in the O-linked glycosylation of mucins, where MUC5B and MUC21 has the potential for target therapy against LUAD. On the other hand, LUSC seems to be associated with a down-regulation of the complement cascade genes and more generally the innate immune response. These events are triggered, in LUSC, by the activation of three key oncogenic pathways, stimulated by p53, cMYC and P-catenin that impair the induction of execution of a local antitumor immune response. Future in-depth studies on the role of these pathways in LUSC may provide interesting opportunities for drug treatments tailored to this challenging lung cancer type.

We also identified and validated in silico a set of genes that can be collectively used as an ensemble to classify LUAD and LUSC in cancer patient samples, through the integration of different but complementary computational techniques. Some of the candidate genes and pathways identified in our study are usual suspects in lung cancer or other cancer types, attesting the validity of our approaches. Moreover, other candidate genes have been poorly investigated and they could entail novel mechanisms in LUAD and LUSC, deserving attention in future investigations.

## 5. Materials and Methods

### 5.1. Pre-processing of RNAseq data from The Cancer Genome Atlas (TCGA)

We downloaded and pre-processed level 3 legacy RNA-Seq data (RSEM count) for LUAD and LUSC with the *GDCquery* of the *TCGAbiolinks* R package [63,64]. The RNA-Seq have been produced using the lllumina HiSeq 2000 mRNA sequencing platform.

We downloaded the data in October 2016 from the Genomic Data Common (GDC) Portal (https://gdc-portal.nci.nih.gov/). An overview of the analyzed samples is reported in **Table** SI. The samples retained for analyses have been pruned by the 19 ‘discordant LUSC’ samples [31] and by samples with low tumor purity (< 60%) according to a consensus measurement of tumor purity [65].

We then employed the *TCGAbiolinks[63]* function *GDCprepare* to obtain a *Summarized Experiment* object [66]. We removed samples outliers with the *TCGAanalyze_Preprocessing* function of *TCGAbiolinks* using a Pearson correlation cutoff of 0.6. We normalized the datasets for GC-content [67] and library size using the *TCGAanalyze_Normalization* function from *TCGAbiolinks*. Lastly, we filtered the normalized RNA-Seq data for low counts across samples using the function *TCGAanalyze_Filtering*. This step removed all transcripts with mean across all the samples less than 0.25 quantile of the mean. The preprocessed and processed datasets are available through our *Github* repository, along with the script to generate them (https://github.com/ELELAB/LUADvsLUSCtcga).

### 5.2. Differential expression analyses of TCGA datasets

Differential expression analyses have been carried out using *edgeR* [68] and *limma* [37]. The analyses were performed using three different pipelines: one is based on *limma-voom* and the other two are *edgeR-*based, where one of them is developed into *TCGAanalyze_DEA* function, originally incorporated into the *TCGAbiolinks* package (called *edgeR-TCGAb* in this study).

In *limma*, the counts were transformed to log2-counts per million (logCPM) with *voom* [69], which made it possible to apply this tool to RNA-Seq read counts where the robust estimation of the mean-variance relationship replaces the lack of data distribution assumption. In *edgeR* pipelines, the *GLM* (Generalized Linear Models) approach was used by which it is possible to include an experimental design with multiple factors.

In our *limma* and *edgeR* pipelines, the design matrix includes: conditions (tumor vs normal), the patient information when a paired dataset is used, and batches for the other datasets *(unpaired* and *all)*. We corrected for the TSS (Tissue Source Site; the center where the samples are collected) as source of batch effect in our *edgeR* and *limma-voom* DEA pipelines. Other sources of bath effects, such as the plates have been tested in the early stages of the project and did not affect the final conclusion. In contrast, *edgeR-TCGAb* implemented a simpler function for DEA which does not take in account neither the batch correction nor the patient information (only for paired dataset). In all our DEA analyses, we defined as a cutoff to retain significant DE genes a log fold change (logFC) >= 1 or <=-1, whereas a cutoff of 0.01 was used for the False Discovery Rate (FDR).

During the analyses, we also tested two variations of the *limma-voom DEA* pipeline: I) using the same design matrix for *voom* and *ImFit* functions and ii) using the entire *voom* object in the steps following the *voom* transformation and not just the log2-transformed data. These adjustments provide a more correct approach to DEA but did not make any difference on our final conclusions. The corresponding scripts are also reported in our *Github* repository. The overlap between the DE genes identified by each pipeline and for each different curation of the dataset have been estimated using the *UpSetR* package [70].

### 5.3 Curation and differential expression analyses of unified GTEx and TCGA LUAD and LUSC datasets

We used the unified dataset integrating the GTEx [25] cohort of healthy samples and TCGA data as provided by the batch-free *Recount2* protocol [24]. We employed the *TCGAquery_Recount2* function of *TCGAbiolinks v2*.*8*, which we recently developed to query the GTEx and TCGA unified dataset for lung cancer. We then filtered the data for tumor purity with a threshold of 60% as we did for the TCGA dataset (see 3.1) and removing the LUSC discordant samples.

Since *recount2* barcodes were updated to the Universally Unique Identifier (UUID), we carried out a conversion of the filtered TCGA barcodes using the *TCGAAutils* package to be able to apply the pre-processing steps mentioned above with *TCGAbiolinks* (see 3.1). The mapping between the TCGA barcodes and the new UUIDs was carried out by extracting the GDC case identifiers. We analyzed 374, 355 and 393 samples for GTEx, LUAD and LUSC, respectively in the unified datasets. After the filtering steps and preparation of the unified datasets for LUAD and LUSC, only protein-coding genes were retained using the *biomaRt* Bioconductor R package [71–73]. We carried out GC-content normalization and quantile filtering as above (see 3.2). We converted the ENSEMBL identifiers into gene names through the information in the *SummarizedExperiment* object. DEA was carried out with the *limma-voom* method according to the pipeline described above (see 3.2).

### 5.4 Soft-clustering analysis along the clinical stages

We performed gene clustering for the LUAD and LUSC datasets *paired* and *all* according to the normal tumor (NT) and four clinical stages of cancer, i.e stages I, II, III and IV using the *Mfuzz* package version 2.36.0 [45]. *Mfuzz* uses a fuzzy c-means algorithm based on the iterative optimization of an objective function to minimize the variation of objects within the clusters. The benefit of this method is that the fuzzy c-means algorithm is more robust with respect to noise and avoids *a priori* pre-filtering of genes [46]. Analyses by the four stages of cancer and NT were executed to visualize longitudinal evolution of the mean expression in LUAD and LUSC clusters

We carried out the clustering by lung cancer type using the datasets *all*. We built a consensus matrix containing all the gene expression for LUAD and LUSC. The 19 discordant LUSC samples were excluded. We collected all barcodes corresponding to NT samples using the *TCGAbiolinks* function *TCGAquery_SampleTypes* for each lung subtype to identify NT samples in the *all* matrices. We mapped the tumor samples to their stages from the LUAD and LUSC clinical datasets using their barcodes. We filtered out the samples with “not reported” stage status. We computed mean gene expression value per tumor stage and NT for LUAD and LUSC. We settled the parameters for fuzzy c-means clustering using the number of clusters = 6. A minimum standard deviation = 0.0 was used as default parameter. The centroid clustering step results from a weighted sum of all cluster members and shows the overall gene expression pattern in each cluster. The membership values indicate how well a gene is represented by its cluster. Low values illustrate a poor representation of gene by the cluster centroid. Large values point to a high correlation of the expression of gene with the cluster centroid. In each cluster, genes are represented with color lines corresponding to their cluster membership m > 0.56. The membership values are color-encoded in the plots generated by *mfuzz*.*plot*. Yellow or green colored lines correspond to genes with low membership value; red and purple colored lines correspond to genes with high membership value.

### 5.5 Pathway enrichment analyses

We used *ReactomePA* version 1.18.1, an R/Bioconductor package for Reactome Pathway Analysis [44]. We employed the *enrichPathway* function of *ReactomePA* to retrieve the enriched pathways given the DE genes list or the genes from the soft-clustering approach and the background genes list i.e. total genes list used for the DEA. An adjusted p-value cutoff of 0.05 was set and the analysis was done by separating the up- and down-regulated genes for each dataset *{all* and *paired)* and lung cancer subtype. In addition, all the gene symbols were converted to their corresponding ENTREZ IDs provided by the *SummarizedExperiment* object (*GDCprepare* function output).

### 5.6 Gene Ontology (GO) enrichment analysis

To identify biological functions in LUAD and LUSC DE gene sets, we carried out a GO classification, which included the following categories: biological process, cellular component and molecular functions [74].

We performed GO functional enrichment analysis for DEGs using the *topGo* R/Bioconductor package. We provided both DE and background genes lists separating up- and down-regulated genes. The GO results for the biological processes were represented in circular plots generated by the *GOplot* R package [75].

### 5.7 Co-expression network analyses

We used the LUAD and LUSC dataset upon filtering and after *voom* transformation (see 3.2) to carry out modular coexpression analyses with the recently Bioconductor/R package *CEMiTool[15*] using the default protocol suggested by the developers. We also performed pathway enrichment analyses and protein-protein network analyses with the pre-built functions of *CEMiTools*. As a reference for protein-protein interaction we used the *Interologous Interaction Database I2D* version 2.9. [76]

### 5.8 Survival analysis

We performed survival analysis using the R package *survival* version 2.41-3. We used cox regression[77] to estimate differences in survival between patients with low and high expression level of our candidate genes. For each cancer type, tumor samples were extracted and separated by gene expression levels according to lower and upper percentile (25th and 75th, respectively). In cases in which the gene expression level of a specific gene in a certain sample is lower than the 25th percentile the corresponding sample is labelled as *low*, whereas it is labeled as *high* if it the gene expression level is greater than the upper percentile. Since for LUAD, there were tumor duplicates (i.e., tumor samples from the same patient), we calculated their mean for the analysis. The clinical data were downloaded using the *GDCquery_clinical* function of *TCGAbiolinks* and only the patients containing information regarding the last follow-up or death time were included in the analysis.

The cox regressions were performed using the *coxph* function. Cox regression allow to take into account additional explanatory variables, such as age at diagnosis, gender and tumor stage. Before performing Cox regression, we tested the proportional assumption using the *cox*.*zph* function and only the genes that satisfy this test were employed (14 and 16 genes for LUAD and LUSC, respectively). We corrected the p-values of each variable using the Benjamini and Hochberg (BH) method [78].

### 5.9 Independent validation in silico of the candidate genes

To validate our candidate genes, we selected two microarray studies that include both LUAD and LUSC samples. The first study contains 139 and 21 samples for LUAD and LUSC, respectively [79]. The second dataset (GSE33532-GEO accession) 10 and 4 samples for LUAD and LUSC, respectively [80]. At first, we converted the probe sets to gene names using the *gconvert* function of the *gProfileR* package[81] (0.6.6 version) and we removed all the non-converted probes. Since multiple probe sets can identify the same gene, we collapsed them to obtain unique matches with the *collapseRows* function implemented in the WGCNA package-1.63 version [16]. We performed the hierarchical clustering (complete method and euclidean distance) and the results was showed in a heatmap (*heatmap*.*2* function in the *gplots* package 3.0.1 version).

## ACKNOWLEDGEMENTS

The authors would like to thank Antonio Colaprico (University of Miami), Francesco Russo (NNF Center for Protein Research, University of Copenhagen) and Vanna Albieri (Danish Cancer Society Research Center) for fruitful discussion and comments. We also would like to thank Matteo Lambrughi for assistance with the illustrations for figure 8.

## Author contributions

Conceptualization: EP; Data curation: ML, EP; Formal Analysis: ML, IP, EP; Funding Acquisition: EP; Investigation: ML, EP; Methodology: ML, TT, MV, MM, IP, EP; Project Administration: EP; Resources: EP; Software: ML, IP, MM, EP; Supervision: EP; Validation: EP, ML; Visualization: ML, IP, EP; Writing-Original Draft Preparation: EP; Writing-Review and Editing: EP with inputs from all the coauthors.

## Availability of data and materials

The datasets generated and/or analyzed during the current study are available in our Github repository: https://github.com/ELELAB/LUADLUSCTCGAcomparison

