## Supplementary Figure S1 for "Different molecular signatures in lung cancer types from integrative bioinformatic analyses of RNASeq data"

Figure S1. We illustrate the results from GO-enrichment analyses on the unique LUAD up-regulated genes where it is possible to appreciate the biological processes that are enhanced in LUAD, including the O-glycan processing as well as cadherins involved in cell-cell adhesion. The results on O-linked glycosylation are confirmed by the pathway enrichment analyses as well (see Table 1 main text).

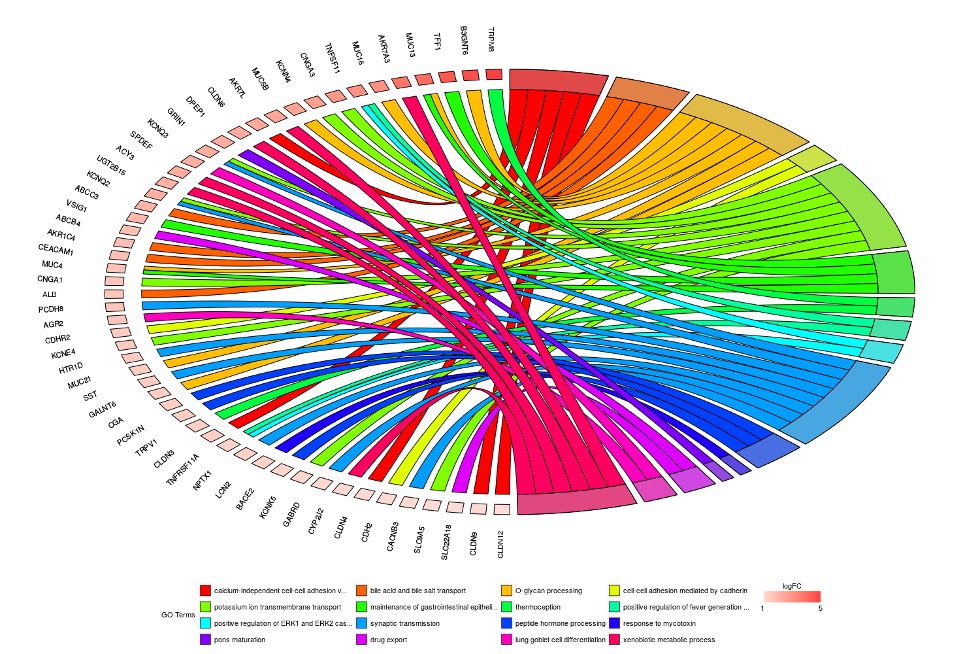
