## Supplementary Figure S2 for "Different molecular signatures in lung cancer types from integrative bioinformatic analyses of RNASeq data"

**Figure S2**. **Soft-clustering across lung cancer clinical stages in LUAD.** Each cluster describes an expression pattern in the dataset through the four stages of cancer i.e stages I, II, III and IV. Blue and purple lines correspond to genes with high cluster membership value, *i.e*. m> 0.56. A Table with the genes belonging to each cluster and their m value is reported in the Github repository. The example of LUAD_all_ is showed here, whereas results for LUSC_all_ are reported in Figure 3.

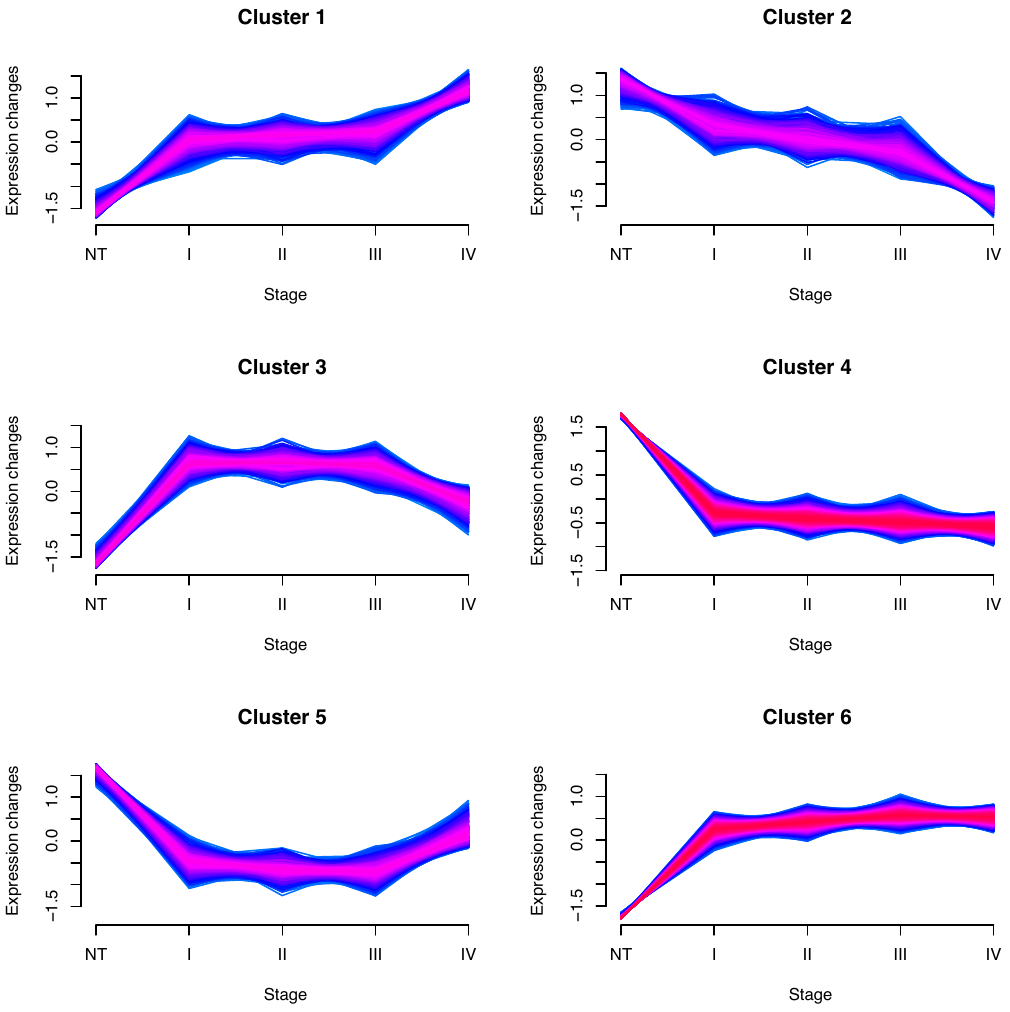
