## Supplementary Table S1 for "Different molecular signatures in lung cancer types from integrative bioinformatic analyses of RNASeq data"

**Table S1. Summary of the datasets used in the study.** NT and TP refers to normal and primary tumor samples, respectively.

|  | NT | TP | TOT mRNA before pre-processing | TOT mRNA after pre-processing |
| --- | --- | --- | --- | --- |
| LUAD_all_ | 59 | 324 | 20330 | 13044 |
| LUAD_unpaired_ | 59 | 292 | 20330 | 13044 |
| LUAD_paired_ | 27 | 32 | 20330 | 13044 |
| LUSC_all_ | 51 | 356 | 20330 | 13044 |
| LUSC_unpaired_ | 51 | 321 | 20330 | 13044 |
| LUSC_paired_ | 35 | 35 | 20330 | 13044 |
