## Supplementary Table S2 for "Different molecular signatures in lung cancer types from integrative bioinformatic analyses of RNASeq data"

**Table S2**. Summary of DEGs detected by different methods and datasets.

|  | **edgeR** | | **limma** | | **edgeR_TCGAb** | |
| --- | --- | --- | --- | --- | --- | --- |
|  | **up** | **down** | **up** | **down** | **up** | **down** |
| ${LUAD}_{all}$ | 2176 | 1306 | 1443 | 1859 | 2316 | 1311 |
| ${LUAD}_{unpaired}$ | 2205 | 1358 | 1455 | 1897 | 2326 | 1353 |
| ${LUAD}_{paired}$ | 1263 | 1927 | 1262 | 1882 | 2111 | 1204 |
| ${LUSC}_{all}$ | 3005 | 2111 | 2506 | 2494 | 3161 | 2021 |
| ${LUSC}_{unpaired}$ | 2963 | 2156 | 2485 | 2481 | 3157 | 2039 |
| ${LUSC}_{paired}$ | 2226 | 2667 | 2221 | 2646 | 2854 | 2143 |
